# Redox Active Plant Phenolic, Acetosyringone, for Electrogenetic Signaling

**DOI:** 10.1101/2023.09.20.558642

**Authors:** Fauziah Rahma Zakaria, Chen-Yu Chen, Jinyang Li, Sally Wang, Gregory F. Payne, William E. Bentley

## Abstract

Redox is a unique, programmable modality capable of bridging communication between biology and electronics. Previous studies have shown that the *E. coli* redox-responsive OxyRS regulon can be re-wired to accept electrochemically generated hydrogen peroxide (H_2_O_2_) as an inducer of gene expression. Here we report that the redox-active phenolic plant signaling molecule acetosyringone (AS) can also induce gene expression from the OxyRS regulon. AS must be oxidized, however, as the reduced state present under normal conditions cannot induce gene expression. Thus, AS serves as a “pro-signaling molecule” that can be activated by its oxidation - in our case by application of oxidizing potential to an electrode. We show that the OxyRS regulon is not induced electrochemically if the imposed electrode potential is in the mid-physiological range. Electronically sliding the applied potential to either oxidative or reductive extremes induces this regulon but through different mechanisms: reduction of O_2_ to form H_2_O_2_ or oxidation of AS. Fundamentally, this work reinforces the emerging concept that redox signaling depends more on molecular activities than molecular structure. From an applications perspective, the creation of an electronically programmed “pro-signal” dramatically expands the toolbox for electronic control of biological responses in microbes, including in complex environments, cell-based materials, and biomanufacturing.

## Introduction

Integrating electronic networks with biological systems generates powerful opportunities to reveal insights about complex natural systems[1] and provides novel methods to confer precise control over such systems[2]. Connecting molecular communication networks to electronics is of emerging interest as it takes advantage of the well-established, simple, modular, and programmable nature of electronic devices. Facilitating electronic communication with biology allows for finely programmed behavior of responsive cells. Owing to the ubiquity of electronic devices, we believe that electrogenetic technologies, where electronics are used to control gene expression, can potentially revolutionize synthetic biology[2,3] and bioelectronic technologies[4,5]. By developing the field of electrogenetics, designing gene circuits with increasingly complex behaviors will become more facile, expanding the capabilities of controlling living cells[6].

A key challenge in connecting biology and electronics lies in their disparate communication modalities: molecular signals carry information based on structure on the cellular level, whereas the flow of electrons governs electronic communication. However, electronic and cellular circuits operate with similar principles and thus could have compatible signal processing[7,8]. Because redox activity is one of biology’s most prevalent signaling modalities[8], redox activity provides a biologically natural way to interconvert between electronic and molecular, cellular systems. Microbes may be well-suited to convert redox signals into cellular pathway signals, as they contain a multitude of sensors that transduce redox signals into structural changes[9]. Electrogenetics may repurpose these redox sensors to control gene expression. For instance, redox-active diffusible molecules, or mediators, can transport electrons between an electrode and cells harboring redox-responsive promoters[10].

Previous demonstrations using redox for electronic control of gene expression rewired redox-responsive regulons such as *soxR*, that is responsive to redox-cycling drugs[11]; *oxyR*, that is responsive to hydrogen peroxide[12]; and KEAP1 and NRF2, responsive to reactive oxygen species[13]. By applying an oxidative potential to mediators ferricyanide and pyocyanin, Tschirhart and others[14] toggled bacterial gene expression that is regulated by transcriptional regulator SoxR. With this system, electronic inputs controlled phenotypes such as swimming[14] and microbial signaling[13], as well as CRISPR-based signal amplification and noise reduction[15]. For a second, distinct mechanism for electrogenetic control, electrode-generated hydrogen peroxide from the reduction of dissolved oxygen[16] was used to electrogenetically modulate the *oxyR* regulon, enabling control of consortia composition[17,18] and improvement of small molecule production in a co-culture[19]. Electrogenetic signaling is a newly emerging field, and the repertoire of electrochemical inducers and their respective genetic parts is limited. Development of new parts for the electrogenetic toolkit, such as redox-linked elements, will be crucial for designing more complex electronics-controlled genetic circuits, increasing the range of specificity, building bio-electronic devices, and connecting to environments rich in redox signaling.

That is, redox signaling is abundant in biology, in contexts including the gut microbiome[20], soil rhizosphere[21], and disease and inflammation[22]. Despite its ubiquitous nature, the mechanisms and networks of redox signaling remain poorly understood[23–25], and the chemical tools for studying redox species often suffer from limitations[26,27]. Electrochemical tools offer a different approach for studying redox, and indeed have already shown promise by revealing new insights, through methods such as mediated electrochemical probing[28,29].

Electronically controlling expression of the *oxyRS* regulon involves the OxyR protein, which in its native state is reduced and inactive. When oxidized by hydrogen peroxide, OxyR undergoes a conformational shift owing to restructuring of disulfide switches[30], binds with RNA polymerase, and positively regulates transcription of its dependent promoters, including the promoters for *oxyS* and *oxyR*[31]. We hypothesized that redox mediators in their oxidized state might also oxidize OxyR, which has a redox potential (E^0^) of -185 millivolts[32]. Here, we investigated redox mediator 1-(4-Hydroxy-3,5-dimethoxyphenyl)ethan-1-one (acetosyringone), a plant-derived phenolic signaling compound[33] (E^0^ = +0.5 volts[34] vs. Ag/AgCl). Like many phenolic compounds, acetosyringone (AS) is produced by plant tissues in response to stress and pathogen infection[35]. The observation that AS contributes to a redox potential burst during the plant’s oxidative response to infection[36] suggests that it acts as a redox-modulating agent affecting pathogen responses. AS was also demonstrated to oxidize thiol groups of thiolated poly(ethylene glycol)[37], providing a putative mechanism by which AS could interact with the thiol groups of the OxyR subunits.

In this work, we demonstrate electronic control of gene expression of the *oxyS* promoter by oxidized acetosyringone, in addition to induction via production of hydrogen peroxide. That is, we show electrogenetic control of *oxyRS* by bolus addition of oxidized acetosyringone to reporter cells and by applying oxidative potential to a mixture of reporter cells and AS. We further demonstrate that distinct electrochemical mechanisms (oxidation of AS at oxidizing potentials, and the reduction reaction for H_2_O_2_ production at reducing potentials) can actuate a single promoter. That is, we show duality in the approach. By applying charge at +0.5 V, one can oxidize AS and subsequently stimulate *oxyRS* gene expression. Transitioning towards negative (reduced) voltages has no effect, then transitioning further to a more reducing voltage (∼ -0.5V) again stimulates expression via the identical promoter. This duality will enable more sophisticated means for electronically programming genetic circuits, potentially including real time measurement and control. Finally, we also show how dynamic application of these transient induction mechanisms can be overlaid to build complex cellular responses. In sum, our findings reveal that oxidized AS is a novel inducer of OxyR, opening up a richer set of possibilities for the prototypical stress-response mechanism and electrogenetics.

## Results

### Characterization of acetosyringone oxidation

We oxidized a 2 mM solution of acetosyringone (AS) in phosphate buffer (PB) by applying a positive potential to the solution in a half-cell setup[38] (**Fig. 1a**). Applying a potential of +0.7 V – higher than the E^0^ of AS – for 0 to 60 minutes was equivalent to application of 0 to -1.74 coulombs of charge. This gradually turned the AS solution from colorless to brownish-orange, indicating its oxidation[39,40] (**Fig. 1b**) and increasing levels of AS oxidation were characterized spectrophotometrically with an increasing absorbance peak at 490 nm (**Fig. 1c**). Interestingly, the color change was linear with applied charge (**Fig. 1d**). AS oxidation was also characterized electrochemically using cyclic voltammetry. When scanning from 0 to 0.7 volts, AS had peak currents at 0.47 V (oxidation peak or E_pa_, peak anodic potential) and 0.417 V (reduction peak or E_pc_, peak cathodic potential), and both peak currents were attenuated with increasing application of oxidative charge (**Fig. 1e**). Again, current attenuation was linear with applied charge (**Fig. 1f**). Oxidation of AS when suspended in phosphate buffer (PB) at pH 7.4 can thus be characterized by absorbance at 490 nm, current at the oxidative peak, and current at the reductive peak, as each of these three metrics correlated to the applied potential.

**Figure 1.**
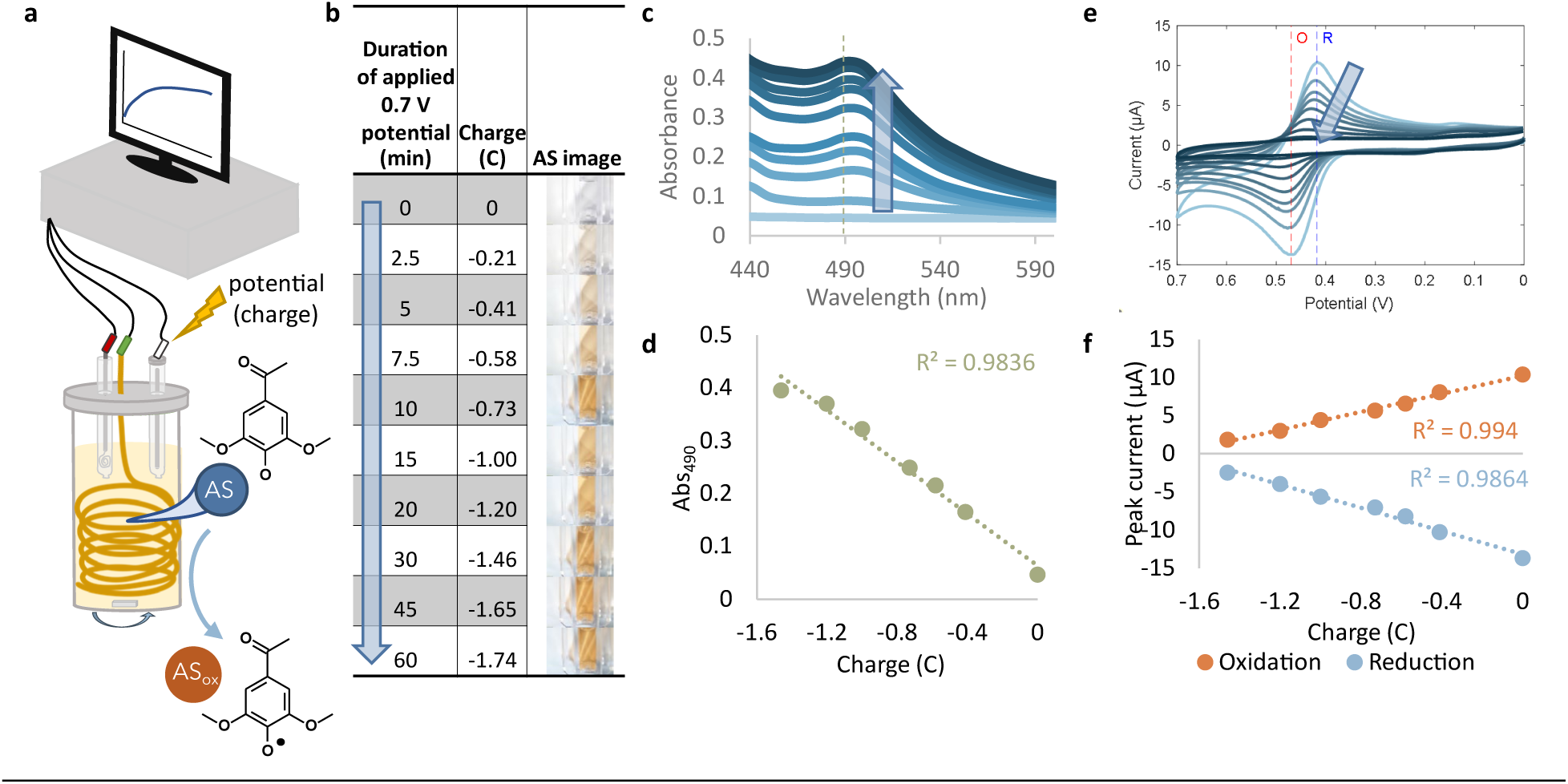
Acetosyringone (AS) electrochemical oxidation is robustly characterized. **(a)** Electrochemical setup for application of charge to AS solution. Gold wire working electrode, Ag/AgCl reference electrode, and platinum counter electrode (separated by a salt bridge) were submerged in AS solution and connected to a potentiostat. Accumulated charge, optical images of AS solution in a cuvette (**b**), absorbance spectra (**c**), and cyclic voltammogram (CV, **e**) of 2 mM AS which was charged (oxidized) with +0.7V for 0-60 minutes. Images are taken immediately post applied charge. Reduction peak (‘R’) and oxidation peak (‘O’) are marked on CV. Arrows indicate application of charge to AS for a longer duration of time. (**d**) Correlation of applied charge to absorbance at 490 nm and (**f**) to currents at oxidation and reduction peaks.

### Oxidized acetosyringone imparts oxidative stress on *E. coli*

Perhaps oxidized AS inhibits pathogen invasion of plants by increasing toxicity towards microbes[41]. While *E. coli* is not a typical plant pathogen, we performed a colony count assay after treating *E. coli* with PB, AS, and oxidized AS (**Fig. 2a**), and we found oxidized AS inhibited *E. coli* growth more strongly than AS in its natural (reduced) state (**Fig. 2b-c**), suggesting a negative impact on the cell’s metabolic state. We hypothesized that the OxyR hydrogen peroxide-responsive stress regulon might be involved in the cellular response to oxidized AS. The OxyR protein is intracellularly oxidized by hydrogen peroxide and activates transcription of genes encoding catalase (*katG*), glutaredoxin (*grxA*), small noncoding transcriptional regulator OxyS RNA (*oxyS*), and other responding molecules to the H_2_O_2_ stress (**Fig. 2d**). Indeed, qPCR revealed that oxidized AS upregulated expression of two OxyR-regulated genes, *oxyR* and *katG* (**Fig. 2e**). Upregulation of genes in the *oxyR* regulon by oxidized AS suggests the potential to use oxidized AS for synthetic gene expression through the OxyR-regulated *oxyS* promoter.

**Figure 2.**
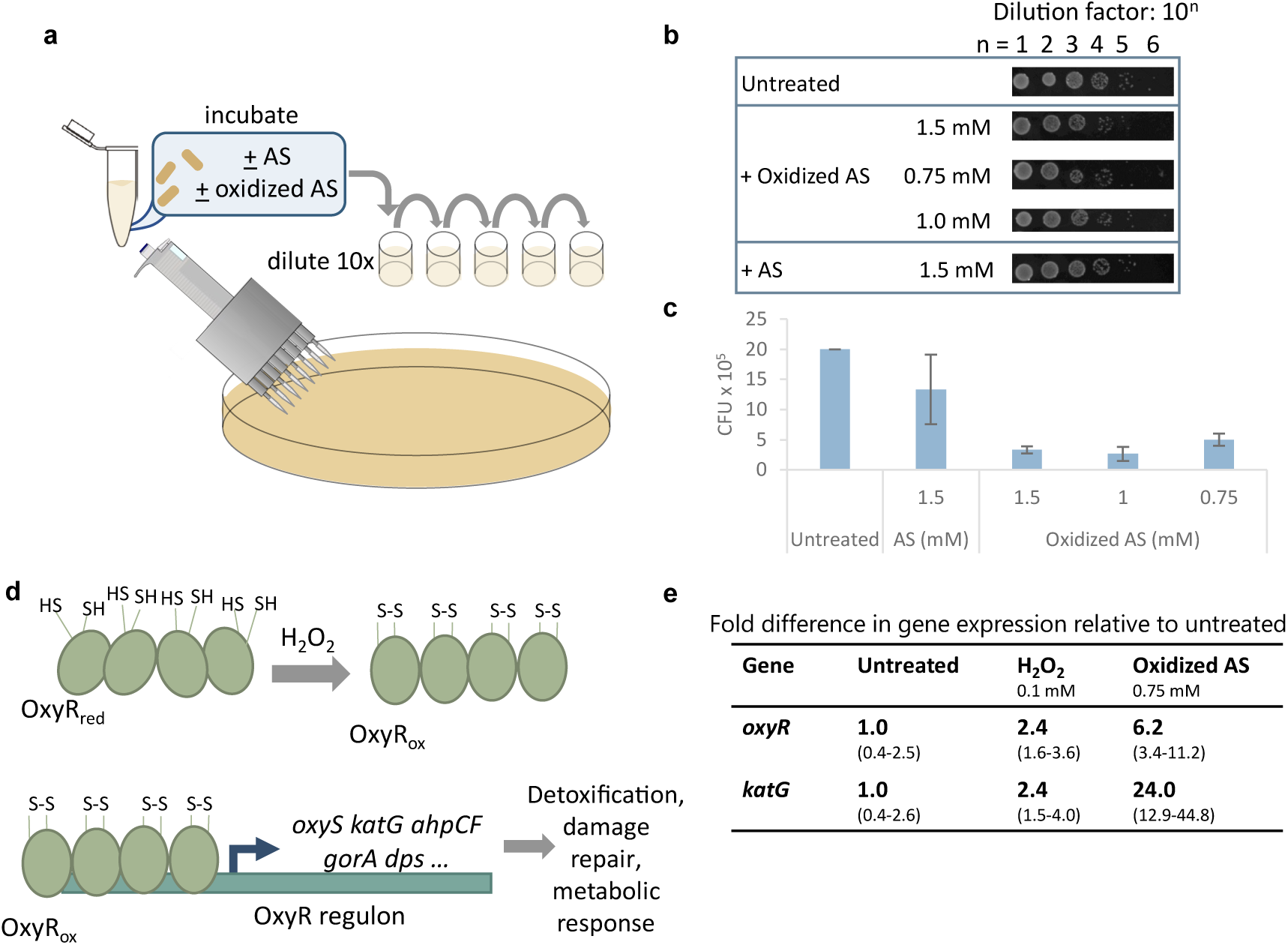
Oxidized acetosyringone elicits cellular oxidative stress response. (**a**) Scheme of colony count assay. (**b**) Colonies and (**c**) colony forming units (CFU) observed after cells were mixed with oxidized and reduced AS. Cells in LB were mixed with AS or phosphate buffer (PB, as a negative control), incubated for 1.5 hours with shaking at 37° C, and plated in 10x dilutions. (**d**) The OxyR regulon in *E. coli*. The OxyR regulator is inactive in its reduced state. Upon oxidation by H_2_O_2_, oxidized OxyR binds to DNA at the promoter region of cognate genes and activates transcription. Adapted from Pomposiello & Demple (2001). (**e**) qPCR data showing effect of oxidized AS on cellular expression of stress-related genes *oxyR* and OxyR-regulated *katG*, with standard deviation of fold difference shown in parenthesis. Cells were grown to OD_600_ = 0.4, treated with the respective inducer, and pelleted at an OD_600_ of 0.8-0.9 for RNA extraction.

### Setting the redox state of acetosyringone enables control of gene expression

To investigate whether oxidized AS could induce expression via hydrogen peroxide-inducible OxyR, we used previously engineered *E. coli* OxyRS-sfGFP reporters which express sfGFP downstream of the *oxyS* promoter[42] (**Fig. 3a**). Applying to AS potentials below its E_pc_ had little effect on OxyRS-sfGFP reporters, while oxidizing AS by applying oxidizing potentials higher than its E_pa_ (0.47 V) induced sfGFP expression resulting in measurable green fluorescence (**Fig. 3b**). The charge accumulated while applying potential to AS served as a gate for induction of the *oxyS* promoter. Applying potentials below the E_pc_ of AS caused minimal accumulation of charge (< -0.086 C), and the resulting AS did not induce gene expression. Applying oxidizing potentials caused charge accumulation over -1.04 C, yielding oxidized acetosyringone which could induce the *oxyS* promoter (**Fig. 3c**). Expression levels were then shown to be modulated by the concentration of oxidized AS and the magnitude of potential applied to AS (**Fig. 3d**), with appreciable fluorescence induced only at higher concentrations and oxidizing potentials (> ∼ 0.5mM, > ∼ 0.5 V). We also found that expression level could be modulated by the duration of the applied oxidative potential (**Fig. 3e**), indicating that both the potential and charge duration applied to AS contribute toward accumulated charge and consequently the extent of AS oxidation. To compare the specificity of oxidized AS as a mediator inducing *oxyR* regulon expression, mediators ferrocene and iridium (E_0_ of +0.25 and +0.67[34] vs Ag/AgCl, respectively) were also tested. Ferrocene and iridium were oxidized by application of +0.3 V or +0.9 V, respectively[34], then added to OxyRS-sfGFP reporter cells in both the reduced and oxidized states. We also measured the final OD_600_ of the cultures to establish effects on cell growth. The two mediators at the concentration ranges tested either did not induce appreciable induction of the reporter cells and had little effect on growth or strongly induced expression but greatly inhibited growth (**Fig. 3f**), suggesting that oxidized AS selectively induces the *oxyS* promoter in a concentration dependent manner and at the same time has minimal effect on growth at concentrations leading to induced expression (i.e. <750 μM).

**Figure 3.**
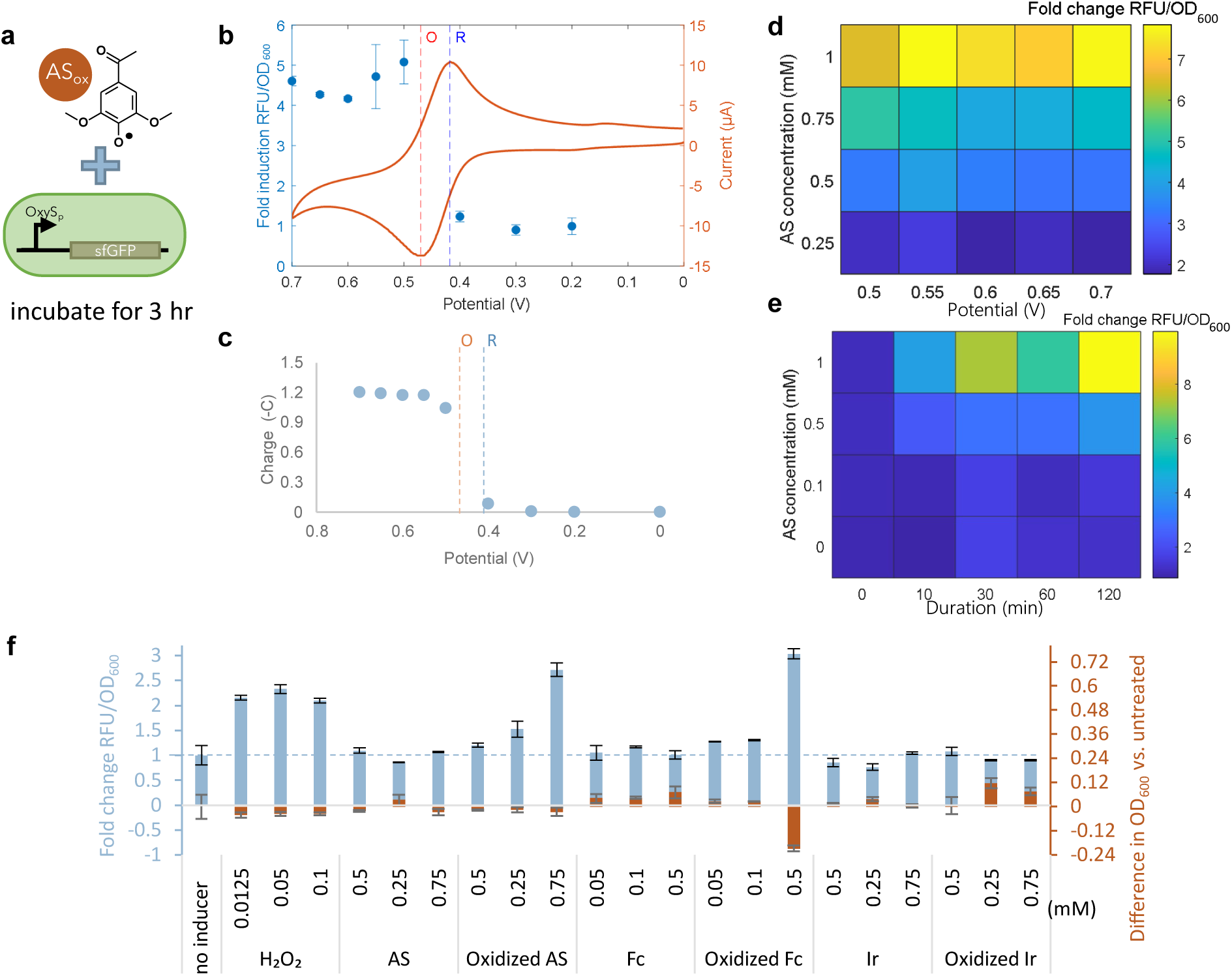
Oxidized acetosyringone (AS) induces the OxyRS promoter. (**a**) Scheme of experiment and gene circuit of OxyR reporter cells. (**b**) The indicated potentials were applied to AS for twenty minutes. OxyR reporter cells were mixed with 0.5 mM AS (to which the potential was applied) and incubated, and fluorescence was measured. Cyclic voltammogram shows oxidative (O) and reductive (R) peaks of AS. (**c**) Accumulated charge after applying indicated potentials to AS for 20 minutes. (**d**) AS was charged at varying potentials for 20 minutes and added to reporter cells at varying concentrations. Reporter cell fluorescence was measured. (**e**) Cell fluorescence resulting from induction by AS (oxidized for varying durations of time at +0.7V) at varying concentrations. (**f**) Fluorescence and final OD_600_ of cells treated with H_2_O_2_, AS, ferrocene (Fc), and iridium (Ir) in reduced and oxidized states. (**b**) - (**f**) show data after three hours of incubation; fluorescence is normalized to OD_600_ and reported as fold change relative to untreated cells.

### Distinct *oxyS* promoter induction dynamics by H2O2 and oxidized acetosyringone

These results show that oxidized AS induces *oxyRS*, and while it is well-known that H_2_O_2_ also induces *oxyRS*, oxidized AS and H_2_O_2_ are electrochemically generated at dramatically different potentials. In **Fig. 4**, we found that both oxidized AS and hydrogen peroxide can induce expression through the OxyR-regulated *oxyS* promoter, but the dynamic responses were quite distinct. Fluorescence from the OxyRS-sfGFP reporter induced by H_2_O_2_ started to rise immediately after addition of H_2_O_2_ and reached a peak 60-80 minutes afterwards (**Fig. 4a**). In contrast, sfGFP fluorescence induced by oxidized AS rose more slowly and leveled off after 70-80 minutes without a subsequent decrease (**Fig. 4b**). The pretreated cells in both cases were identically cultured hence similarly metabolically active. The different responses to H_2_O_2_ and oxidized AS were further highlighted by using OxyRS-sfGFP-AAV reporter cells in which sfGFP was fused with an ssRA[43] degradation tag on its C-terminus. In this way, the sfGFP measurement is more reflective of the rate of generation as opposed to the absolute level. H_2_O_2_-induced fluorescence in the OxyRS-sfGFP-AAV reporters rose rapidly immediately after H_2_O_2_ addition, peaked after 20 minutes, and subsequently fell to the earlier uninduced reporter cell levels (**Fig. 4c**). These data indicated that the response to hydrogen peroxide addition was strong and finite in time, likely owing to the dissolution of inducer, H_2_O_2_, at these cell densities[44]. Oxidized AS, on the other hand, induced a slower rise and a more sustained fluorescence (**Fig. 4d**), indicating continued oxidative stress (and expression) throughout.

**Figure 4.**
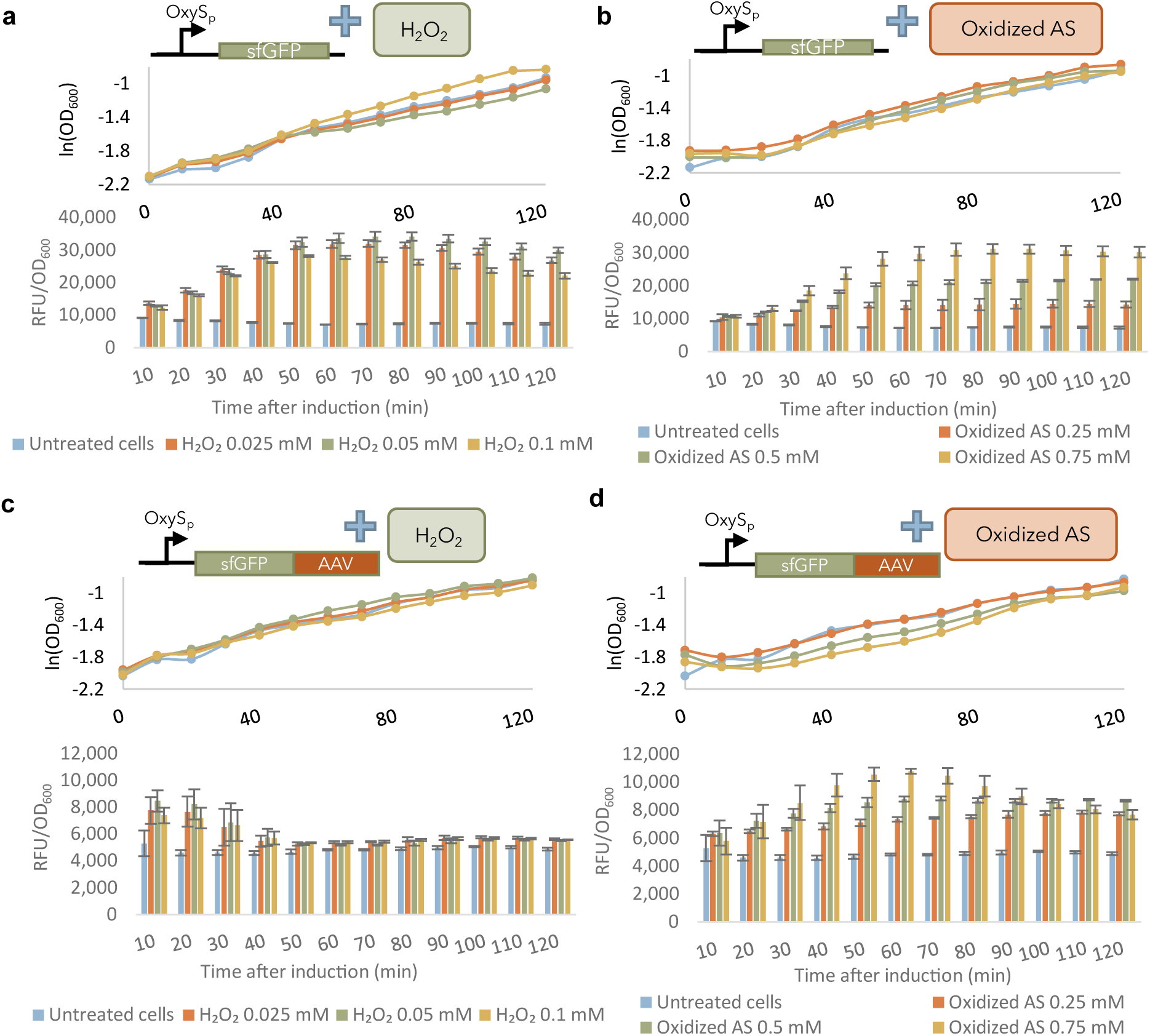
Oxidized AS elicits a distinct response from the OxyRS promoter. OD_600_ and fluorescence of cell reporters induced with H_2_O_2_ or with AS oxidized for 20 minutes at +0.7V. Cell reporters expressed sfGFP under control of the OxyRS promoter with (**a**-**b**) or without (**c**-**d**) a degradation tag (AAV). (**a**)-(**d**) show fluorescence normalized to OD_600_.

We found minimal differences in growth rate between the different induction methods at the selected H_2_O_2_ and oxidized AS concentrations. Further, we performed a simple dynamic analysis of this system, calculating a zeroth order maximum rate expression and a first-order GFP decay rate spanning the appropriate times in **Fig. 4a-d** (**Supplementary Fig. S1** and **Supplementary Table S1**). We found the maximum rate of synthesis was nearly identical for all hydrogen peroxide addition cases without a degradation tag (**Fig. 4a**) and the oxidized AS case at the 0.75 mM level (**Fig. 4b**). Interestingly, we also found nearly linear dependence of this maximum synthesis rate with AS concentration. We next found that the first-order degradation rate for the H_2_O_2_-induced sfGFP-AAV case (with the degradation tag) was roughly 3-fold larger than the largest degradation rate for the AS-induced sfGFP-AAV case (**Supplementary** Fig. 1 and **Supplementary Table S1**). This was somewhat surprising and suggested that the response to hydrogen peroxide elicited more proteolytic activity than the AS addition, although this was not tested. These results do show, however, that the responses of the cells to the different methods of induction were different.

### Electronic control of gene expression

A promising outlook in electrogenetic control is to turn gene expression “on” by simple application of an electronic cue[14]. To demonstrate, we added AS to reporter cells and subsequently applied an oxidizing potential. This differs from the oxidized AS induction described in the previous experiments, where a bolus addition of oxidized AS induced fluorescence. Instead, here we show direct electrogenetic control: the signal for inducing expression of the OxyRS-sfGFP reporter cells is the application of an oxidative potential to an electrode immersed in a growing cell culture, rather than bolus addition of an oxidized chemical inducer (**Fig. 5a**). The electrochemical half-cell setup was similar to the setup for oxidation of AS, but we used a 1:1 mix of reporter cell culture and AS diluted in PB to the desired concentration (**Fig. 5b**). When an oxidizing potential was applied to the mixture of cell culture and AS, an increase in fluorescence was observed. This was directly analogous to the results from bolus addition of oxidized AS to reporter cells (**Fig. 5c** vs **Fig. 4b**). Gene expression and fluorescence could be tuned by varying the concentration of AS added as well as the duration of the applied oxidative voltage (**Fig. 5d**). Higher AS concentrations and application of oxidizing potential for a longer duration increasingly inhibited cell growth (**Fig. 5e**), such that inducing reporter cells using this method requires optimizing AS concentration and applied charge to avoid altered cell growth. Varying the potential applied also provides significant power to modulate gene expression (**Fig. 5f**). Importantly, we found that the accumulated applied charge correlated linearly with GFP fluorescence in all cases, whether the parameter being varied was charge duration or potential (**Fig. 5g**). That is, we show here that gene expression can be reliably tuned based on total accumulated charge, and either potential or charge duration can be adjusted for fine-tuned modulation of gene expression.

**Figure 5.**
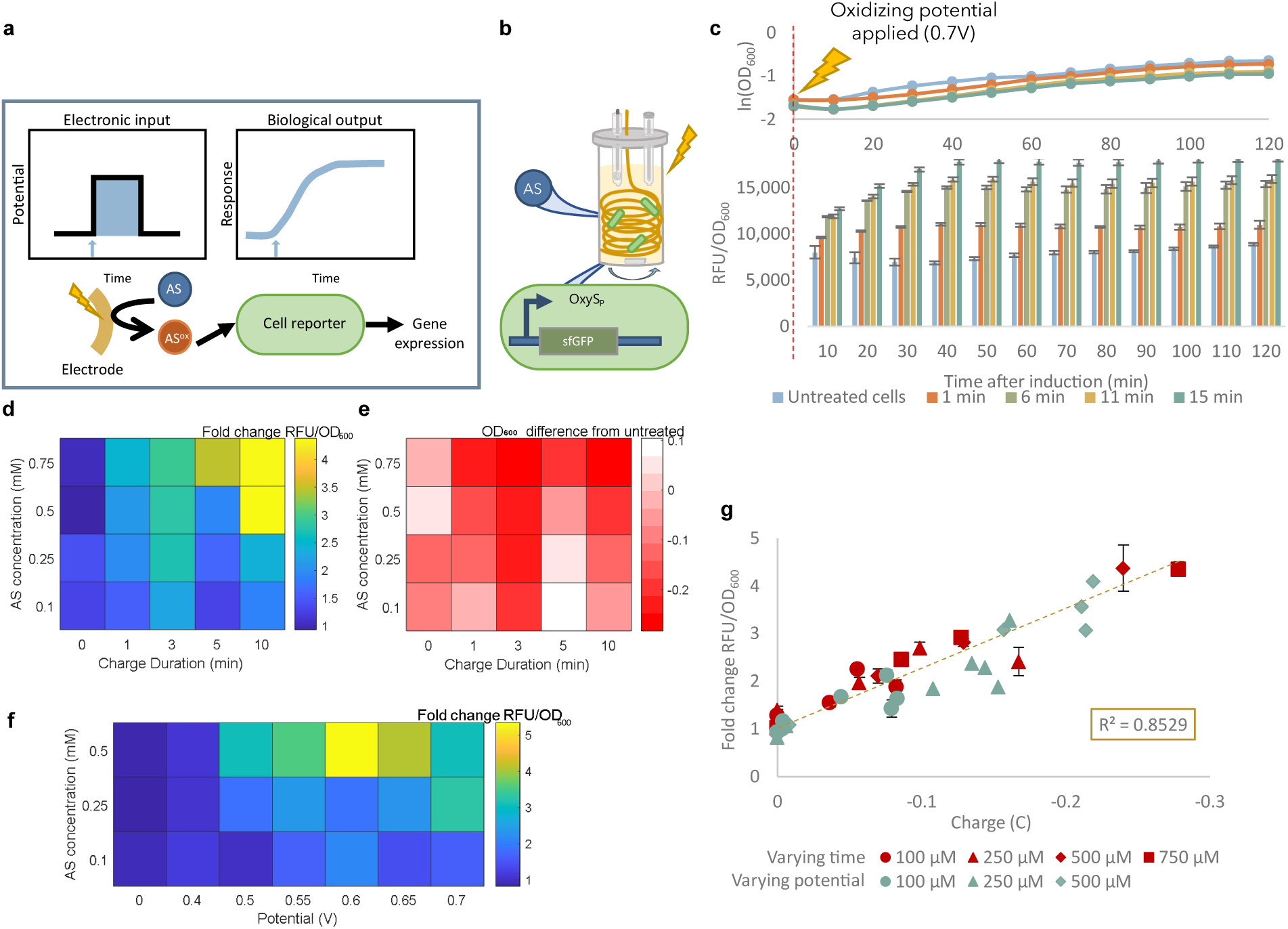
Acetosyringone (AS) can be applied for electronic control of cell fluorescence. (**a**) Direct electrogenetic control scheme. A potential applied through the gold electrode causes oxidation of AS in solution, which induces a cellular response in the cell reporters via the *oxyS* promoter. (**b**) Scheme of the electrochemical setup. A gold wire working electrode, Ag/AgCl reference electrode, and platinum counter (separated by a salt bridge) are submerged in a 2.4 mL mixture of liquid cell culture and AS. (**c**) OD_600_ and fluorescence of cell reporters induced by direct electrogenetic control. 100 μM AS was added to reporter cells, and an oxidative potential of +0.7V was applied for the indicated duration. Fluorescence is reported as the moving average of fluorescence over 30 minutes with 3 technical replicates. (**d**) Fluorescence of reporter cells after addition of AS at varying concentrations and application of +0.7V for varying durations of time. Fluorescence was normalized to that of untreated cells, and (**e**) the difference in OD_600_ between treated and untreated cells three hours after induction is shown. (**f**) Fluorescence of reporter cells, normalized to untreated, after addition of AS at varying concentrations and application of varying potentials for 7.5 minutes. (**g**) Correlation of applied charge to cell fluorescence. Cells were mixed with the indicated concentrations of AS. A constant potential of +0.7V was applied for varying durations from 0-10 minutes (red), or the potential was varied from 0 to +0.7V and applied for twenty minutes (green). The linear regression line, calculated for all values whether varying time or varying potential, and R^2^ value are shown. (**c**), (**f**), and (**g**) show fluorescence after three hours of incubation normalized to OD_600_ and are reported as fold change relative to untreated cells.

### An electronic switch: a duality in controlling gene expression

It is important to note that electrogenetic control of the *oxyS* promoter is also achieved using the 2-electron oxygen-reduction reaction (ORR) to convert dissolved oxygen into H_2_O_2_, thereby inducing OxyR-regulated gene expression by electronically generating one of its native signals[15,17,38]. The ORR is carried out at reducing potentials (-0.5 V with a gold electrode [18,38]) whereas oxidation of AS occurs at oxidizing potentials, and both H_2_O_2_ and oxidized AS are now shown to induce expression via OxyR (**Fig. 6a)**. To verify induction of OxyRS-sfGFP reporters by electrode-generated H_2_O_2_ and to explore the extent to which one can use varied inputs, we applied potentials varying from +0.8 to -0.7 V to reporter cell cultures in the absence of any mediator. As expected, only application of the reducing potentials required for ORR (from -0.5 to -0.7 V[38]) accumulated charge (**Fig. 6b**), consistent with generation of H_2_O_2_ at the electrode[38]. Interestingly, when the same experiment was repeated in the presence of acetosyringone, charge accumulation was observed both at oxidizing potentials (+0.5 to +0.8 V) and reducing potentials (-0.5 to -0.7 V, **Fig. 6b**). While most of the applied potentials resulted in comparable growth rates and final OD_600_ values, applying oxidizing potentials inhibited growth only in the presence of AS, consistent with earlier observations (**Fig. 6c**). Gene expression followed the same trends as the accumulated charge. Without AS present, only reducing potentials elicited fluorescence (“reductive activation”), whereas when AS was present, fluorescence was observed at both reducing and oxidizing potentials (“oxidative activation”) (**Fig. 6d** and **Supplementary Fig. S2**). Interestingly, induction by oxidation of AS led to higher fluorescence per unit charge (**Fig. 6e**). These results suggest that AS can facilitate gene expression at two distinct potential ranges: at oxidizing potentials, as oxidized AS induces the *oxyS* promoter; and at reducing potentials, as the presence of AS does not interfere with ORR-based H_2_O_2_ generation, allowing for induction of the *oxyS* promoter by H_2_O_2_.

**Figure 6.**
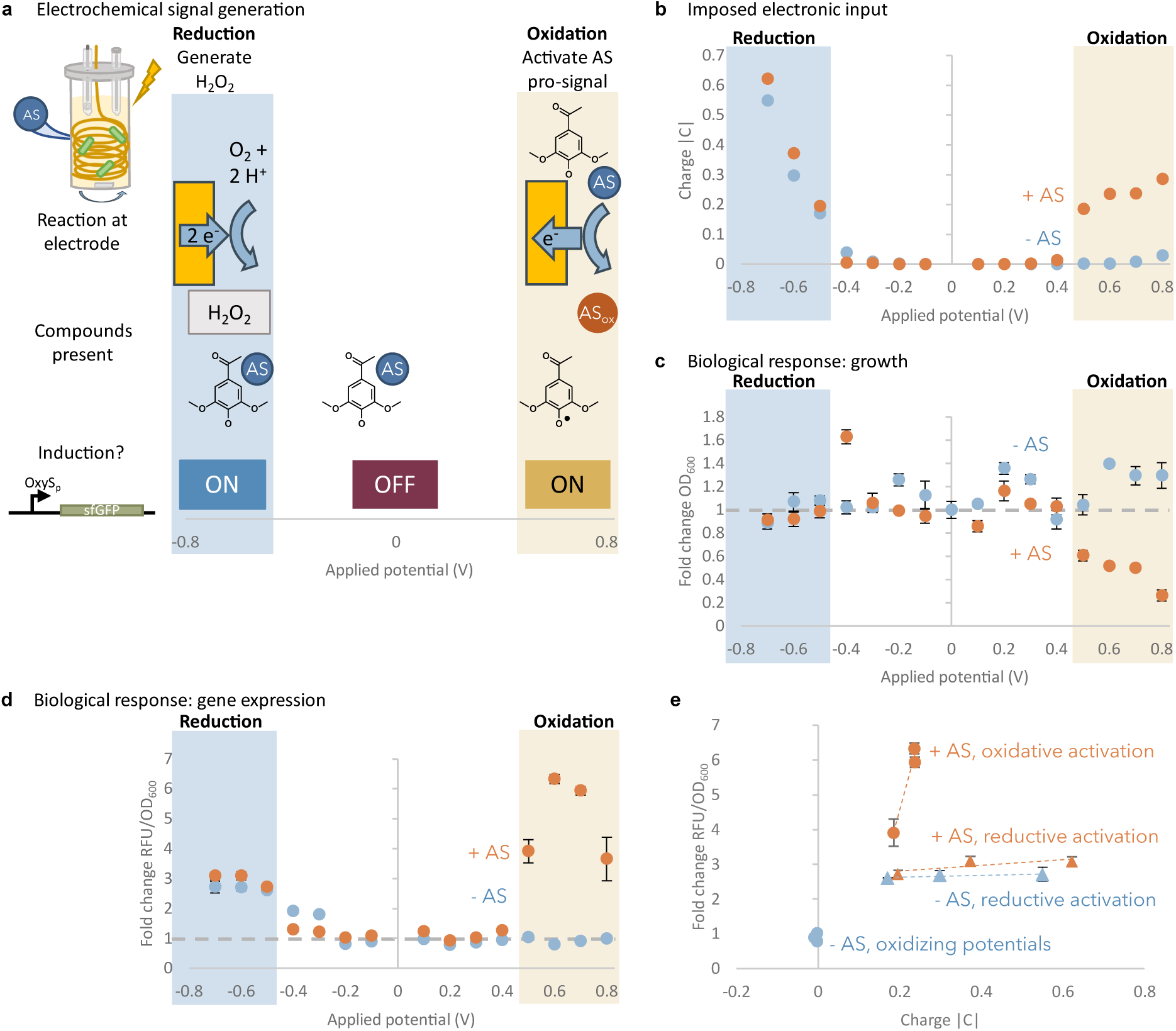
Biological responses (growth and gene expression) are elicited by electrochemical signals generated at distinct potential ranges. (**a**) Result of applying electronic signals to cells in the presence of AS. Application of a reducing potential leads to H_2_O_2_ generation due to the oxidation-reduction reaction. Application of an oxidizing potential generates oxidized AS. Both H_2_O_2_ and oxidized AS induce gene expression via the OxyR regulon. (**b**) Cumulative applied charge, (**c**) OD_600_ after three hours of incubation normalize to untreated cells, and (**d**) fold fluorescence of POxyRS-sfGFP reporter cells after varying potentials are applied for 7.5 minutes in the absence or presence of 500 μM AS. Fluorescence is reported after three hours of incubation and is normalized to OD_600_, then the RFU/OD_600_ value is normalized to that of untreated cells. (**e**) Correlation of cumulative applied charge to gene expression for cells to which AS was added or omitted and reducing (-0.5 to -0.7) or oxidizing (+0.5 to +0.7) potentials were applied. Linear regressions have a slope of 0.25 (-AS, reducing potentials), 0.81 (+ AS, oxidizing potentials), and 43.0 (+ AS, oxidizing potentials).

### Dynamic electrogenetic programming

To compare our acetosyringone-based electrogenetic approach with ORR-based H_2_O_2_ production, we mixed acetosyringone with OxyRS-sfGFP-AAV reporter cells and treated in varied electronic modalities by applying 5-minute “pulses” of either oxidizing potential (+0.7V) or reducing potential (-0.5 V) and different combinations thereof (**Fig. 7a**). In **Fig. 7b**, we pulsed +0.7V in a repeated fashion for 300 seconds (-160 ±47 mC charge) every 45 minutes. It was readily evident that the added oxidizing potential repeatedly induced the cells, and interestingly, nearly equivalently at each step. The degradation-tagged sfGFP showed peaks in fluorescence followed by decreasing levels starting from 30-90 min after the pulse, back towards a still induced but lower level. With multiple applications of a reducing potential (-0.5 V for 300 seconds, or +191 ±55 mC charge), we found sharp peaks followed by more rapid decay until all reached similar levels typically an hour after the peak in fluorescence (**Fig. 7c**). Subsequent tests with alternating oxidation and reduction pulses resulted in varied responses with different patterns, peaks, and decay times (**Fig. 7d-7f**). Interestingly, in each case where +0.7 V was applied, there was a relatively slower response (in both increase and decrease of signal), while in each case where -0.5V was applied, there was a faster, sharper increase in fluorescence (**Fig. 7b-7f**). Also, we note that typically, the resultant steady state values were apparently additive, such that a culture with three pulses reached higher levels than those with two and these were seen higher than those with a single pulse, even without normalizing expression to OD_600_ (**Supplementary Fig. S3**). Hence, applying multiple pulses to the reporter cell culture over time yielded somewhat predictable and additive fluorescence outputs, demonstrating dynamic temporal control of gene expression based on the type (potential), duration, and timing of the electronic input.

**Figure 7.**
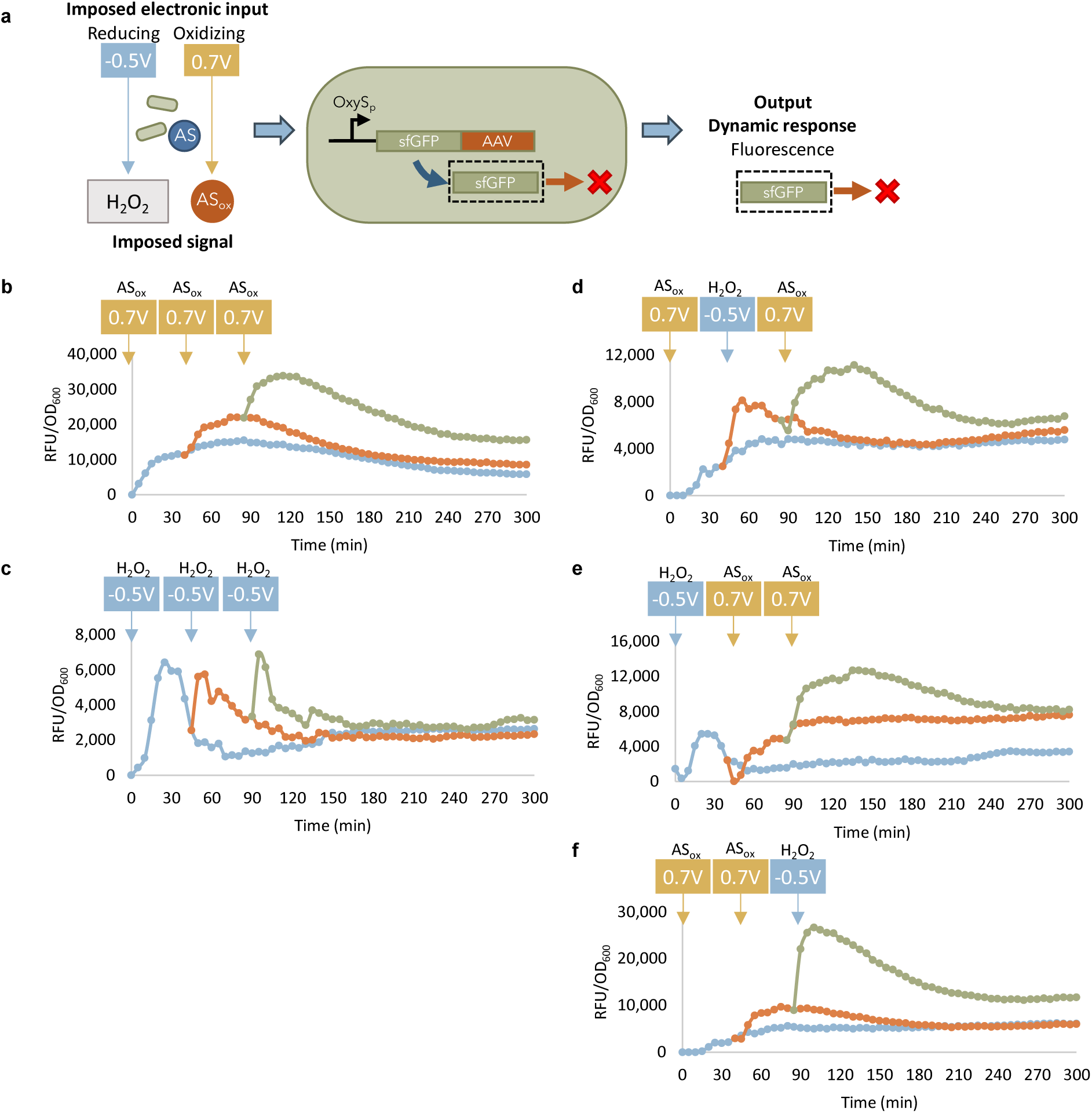
Electrochemically generated signals influence gene expression dynamics. (**a**) Scheme of electrochemical signals generated at reducing and oxidizing potentials in the presence of AS and reporter cells expressing degradation-tagged fluorophores. Fluorescence of POxyRS-sfGFP-AAV reporter cells, in the presence of 500 μM AS, after application of “pulses” of potential at the indicated voltage and time for 7.5 minutes. If the value of fluorescence difference between the sample and the untreated cell negative control was less than 5 RFU, it was denoted as 0 RFU. The applied potential “pulses” were (**b**) three oxidizing pulses, (**c**) three reducing pulses, (**d**) one reducing followed by two oxidizing pulses, (**e**) an oxidizing, reducing, then oxidizing pulse, or (**f**) two oxidizing pulses followed by a reducing pulse.

## Discussion

Electronic control of gene expression expands our ability to program and manipulate cellular behavior. In this work, we demonstrate that redox electrogenetics can be facilitated by acetosyringone via transcriptional regulator OxyR and the *oxyS* promoter. OxyR has been described to be primarily activated by hydrogen peroxide and partially activated by diamide and S-nitrosocysteine[32]. Oxidized acetosyringone, a phenolic plant-produced redox mediator, is reported here for the first time to be an inducer of OxyR. Importantly, this work introduces a new mechanism and electronic potential range for tunable electronic control of gene expression. We show that OxyR-mediated expression can be electronically induced both at reducing potentials (< -0.5 V[17]) by electrode-generated H_2_O_2_, and at oxidative potentials (> 0.7 V) by oxidized acetosyringone (**Fig. 6a**). Induction of SoxR-mediated gene expression has previously been achieved at oxidative potentials only (> 0.3 V[14]), and the electrochemical configuration was based on two mediators - pyocyanin and ferricyanide/ferrocyanide - as well as the maintenance of an anaerobic environment. Acetosyringone-facilitated induction, on the other hand, provides a simple, tunable, and accessible electrogenetic scheme based on a single mediator that works in an aerobic environment, thus expanding the range over which electronic activation can be applied. In addition, because each of the described methods for electronic induction are accessed at distinct potential ranges (more negative than -0.5 V, more positive than 0.3 V, more positive than 0.7 V), future work can multiplex electronic inputs by utilizing applied potential as a “knob” to selectively toggle expression of desired genes.

To our knowledge, this is the first demonstration harnessing the redox signaling capabilities of acetosyringone for electronically actuating cellular function. Acetosyringone is a natural mediator produced by plants during wound stress and has been studied as a signaling molecule that reportedly conveys information by nature of its molecular structure. Specifically, acetosyringone induces chemotaxis and virulence gene expression in *Agrobacterium* through the *vir* regulon[45]. This work now shows that that AS can also act as a signaling molecule through a distinct mechanism - by virtue of its redox properties. It is silent towards OxyR until “activated” by an oxidative potential. In this capacity, AS serves as a “pro-signal”, wherein its messaging is conveyed when it is in the oxidized form. “Pro-signals” are consistent with the concept that redox signaling depends more on molecular activities than molecular structure[8], as the OxyR transcription factor appears to respond similarly to both H_2_O_2_ and oxidized AS. Oxidized acetosyringone has been shown previously to elicit stress responses in *Pseudomonas syringae*, upregulating expression of genes involved in metabolism, energy generation, and cell wall components and inducing a viable but not culturable (VBNC) state[46]. Our work builds upon such insights by highlighting a particular cellular response elicited specifically by oxidized AS but not reduced AS. As OxyR and hydrogen peroxide-responsive homologs are widely conserved across many bacteria[31], future applications of this work may involve sensing or actuation of more rhizosphere-relevant bacteria, perhaps in response to an oxidative AS signal released by an infected plant.

Interestingly, we observed a difference in the dynamics of the OxyR-regulated responses elicited by H_2_O_2_ and oxidized AS. Importantly, we show that a single promoter can be toggled using two distinct mechanisms simply by varying the type of potential applied to cells in the presence of AS. The imposed electronic input, or redox context, defines the duration and magnitude of the biological response, thus further demonstrating the importance of context for redox activity[47,48]. Both H_2_O_2_ and oxidized AS are signaling molecules present in the rhizosphere after plant infection. The difference in the dynamic responses to the two signaling molecules may have implications relative to the bacterial response to plant-produced oxidative stressors.

By selectively tuning and applying oxidizing and reducing potentials in pulses, we can generate patterns of oscillating gene expression (**Fig. 7**). Oscillatory patterns are natively found in biology, including signaling molecules that are expressed in pulses[49] and plant responses to multiple stresses[50]. As such, our electrochemical setup using acetosyringone can be used to recapitulate natural oscillatory activity. At the same time, biomanufacturing applications could involve synthetic strains to improve production of relevant molecules[51], or activate a particular cellular response that is present for either a transient period or for a sustained duration depending on whether the inducer present is H_2_O_2_ or oxidized AS.

Understanding the signaling characteristics of oxidized acetosyringone provides new avenues for learning and building technologies for the soil rhizosphere as well as for engineered systems. Using the acetosyringone induction system, one could envision biosensors, cells and cell networks that collect new types of information about their local redox state wherein they are programmed to carry out designer functions (e.g., generate nutrients for root structures, degrade recalcitrant contaminants, or otherwise synthesize value-added products).

## Materials and Methods

### Strains and plasmids

The strains, plasmids, and primers used in this work are listed in **Table 1**. Plasmid pOxyRS-sfGFP-AAV is derived from pOxyRS-sfGFP[37] and incorporates a *ssrA* degradation tag encoding AANDENYLAAAV[43] (“AAV”) at the 3’ end of sfGFP.

**Table 1.**
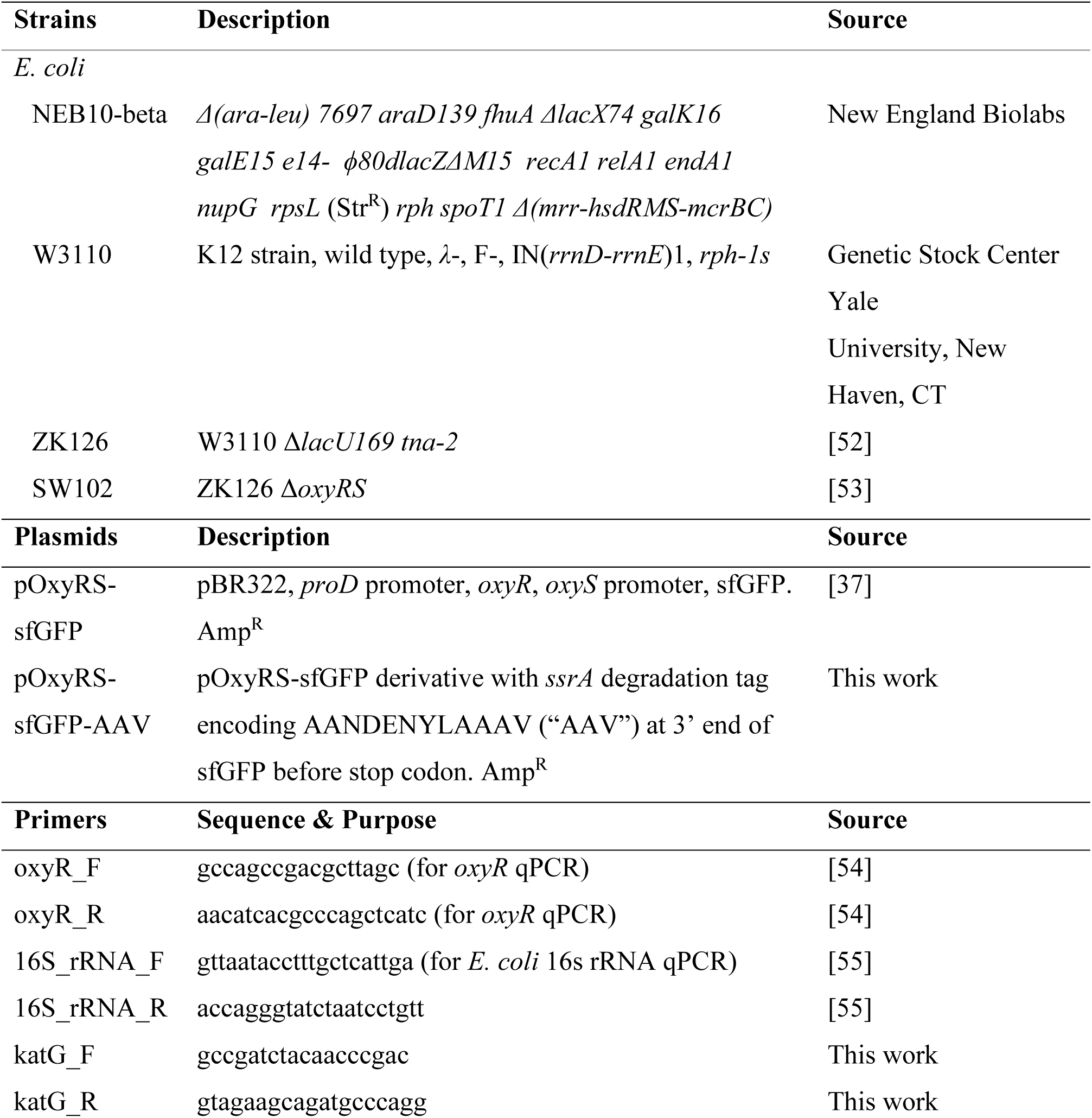
Strains, plasmids, and primers used in this study.

### Cell culture

Plasmid pOxyRS-sfGFP was cloned into strain SW102 (ZK126 ΔOxyRS) and plasmid pOxyRS-sfgFP-AAV was cloned into NEB10-beta. Cells were cultured at 37 °C with shaking at 250 rpm in an incubator or in a TECAN SPARK microplate reader. LB media was used for cloning and for growing 4-5 mL overnight cultures. For cell experiments, overnight cultures were diluted 100x into M9 media (1x M9 salts, 0.1 mM CaCl_2_, 2 mM MgSO_4_, 0.4% glucose, and 0.2% casamino acids) with the appropriate antibiotics and cultured until the OD_600_ reached ∼0.4 before addition of inducer(s). Unless indicated, all other cell experiments used a 1:1 mixture of M9 media and 0.1 M phosphate buffer (PB, pH = 7.2) (with acetosyringone when indicated). For ampicillin-resistant plasmids (pOxyRS-sfGFP and pOxyRS-sfGFP-AAV), ampicillin was used at a concentration of 100 μg/mL. Microplate reader experiments were conducted using 96-well plates and 200 μL sample volume, and samples were measured in triplicate.

### Electrochemical setup

Electrochemical techniques were run using a CHI 6273 C electrochemical analyzer (CH Instruments). All electrochemical methods used a three-electrode setup in a 17 mm diameter glass vial with an Ag/AgCl reference electrode, platinum wire counter electrode, gold working electrode, and small stir bar. For cyclic voltammetry, a 2 mm gold standard working electrode (CH Instruments) was used, and potential was scanned from 0 to 0.7 V and back at a scan rate of 0.2 V/sec with a 0.001 V sample interval and 1x 10^-5^ A/V sensitivity. For routine oxidation of acetosyringone with a half-cell setup[38], a 4.5 mL, 2 mM solution of AS (Sigma-Aldrich) in PB was prepared, and the electrochemical setup used a salt bridge as well as a 0.5 mm diameter gold wire (Sigma-Aldrich) approximately 75.4 cm long, coiled to fit in the vial. Using the amperometric i-t curve technique, a constant potential of 0.7 V was applied for 20 minutes with a 0.1 second sample interval and 1x 10^-3^ A/V sensitivity. The potential and duration of applied charge were varied as described in the experimental section. For oxidation of cell culture, the same setup was used, but with 2.4 mL of a 1:1 mixture of cell culture (OD_600_ of 0.4) and either PB only or AS and PB. AS concentration, potential, and duration of applied charge were varied as described in the experimental section. For application of “pulses” of potential, a 2.4 mL of a 1:1 mixture of cell culture (OD_600_ of 0.4) and PB with AS (final AS concentration of 500 μM) was prepared and the indicated potential was applied for 7.5 minutes. 50 μL samples were extricated in triplicate, diluted to 200 μL with equal volumes of M9 media and PB, and placed in a 96-well plate for incubation and measurement in the plate reader. Simultaneously, the remaining cell solution was incubated in a shaking incubator at the same temperature and shaking speed. After 45 minutes of incubation, the cell solution was replenished with 150 μL of a mixture of M9, PB, and AS to the same concentration as the initial setup. The same steps were repeated for a second and third “pulse” of potential.

### Spectrophotometric and fluorometric readings

Absorbance and fluorescence were measured using a TECAN Spark microplate reader. Absorbance spectra of AS were measured with a clear 96-well plate. For cell experiments, 200 μL samples were loaded in triplicate in a black 96-well plate with a clear bottom. The plate was placed in a humidity cassette in the microplate reader, set at 37° C with shaking, and OD_600_ and sfGFP fluorescence (485 nm excitation and 520 nm emission) were continually measured. Fluorescence (relative fluorescence units, or RFU) and absorbance units were normalized by subtracting measurements of the PB/M9 blank sample. Fluorescence was further normalized by dividing by OD_600_ (RFU/OD_600_); when indicated, RFU/OD_600_ was normalized to the RFU/OD_600_ of untreated cell samples (Fold change relative to untreated).

### qPCR assay

6 mL of NEB10β cells were grown to an OD_600_ of 0.4 and induced with PB (negative control), 100 μM H_2_O_2_, or 0.75 mM of oxidized AS. After 30 minutes incubation, 1 mL of cells was pelleted. RNA was isolated with the TRIzol® Max™ Bacterial RNA Isolation Kit (ThermoFisher) according to the manufacturer’s protocol and treated with DNAse I (New England Biolabs). Quantitative real time PCR was run on Applied Biosystems QuantStudio 7 Flex (ThermoFisher) using ∼200 ng RNA, 1.5 μM of the qPCR primers listed in **Table 1**, and the Power SYBR® Green PCR RT-PCR mix following the manufacturer’s protocol (ThermoFisher). Gene expression fold change averages and standard deviations were calculated using ΔΔCT relative to *E. coli* 16s rRNA as an internal control and compared to the negative control of untreated cells.

### CFU count assay

250 μL of NEB10β cells grown to an OD_600_ of 0.4 were transferred to microcentrifuge tubes and treated with PB (negative control), AS, or oxidized AS. Tubes were incubated for 1 hr at 37° C, then four 10x serial dilutions were prepared in PB. 5 μL of each dilution was pipetted onto an LB agar plate in triplicate and incubated overnight. Individual colonies were counted at the highest dilution and colony forming units (CFU) were determined based on the dilution factor.

## Acknowledgements

The authors would like to acknowledge partial support of this work by the National Science Foundation (MBC# 2227598, CBET# 1932963), the Department of Energy (BER#SCW1710), the Defense Threat Reduction Agency (HDTRA1-19-1-0021), and the Gordon and Betty Moore Foundation (#11395). This material is based upon work partially supported by the National Science Foundation Graduate Research Fellowship Program under Grant No. DGE 1840340. Any opinions, findings, and conclusions or recommendations expressed in this material are those of the author(s) and do not necessarily reflect the views of the National Science Foundation.

## Author contributions statement

FRZ wrote the main manuscript text and prepared figures. All authors reviewed the manuscript.

## Data Availability Statement

The datasets generated during and/or analyzed during the current study are available from the corresponding author on reasonable request.

