## Supplementary Information for "Redox Active Plant Phenolic, Acetosyringone, for Electrogenetic Signaling"

[1]Fischell Department of Bioengineering, University of Maryland, College Park, Maryland, United States

[2]Institute for Bioscience and Biotechnology Research, Rockville, Maryland, United States

[3]Robert E. Fischell Institute for Biomedical Devices, University of Maryland, College Park, Maryland, United States

[4]Division of Biology and Biological Engineering, California Institute of Technology, Pasadena, California, United States

### Supplementary Figures

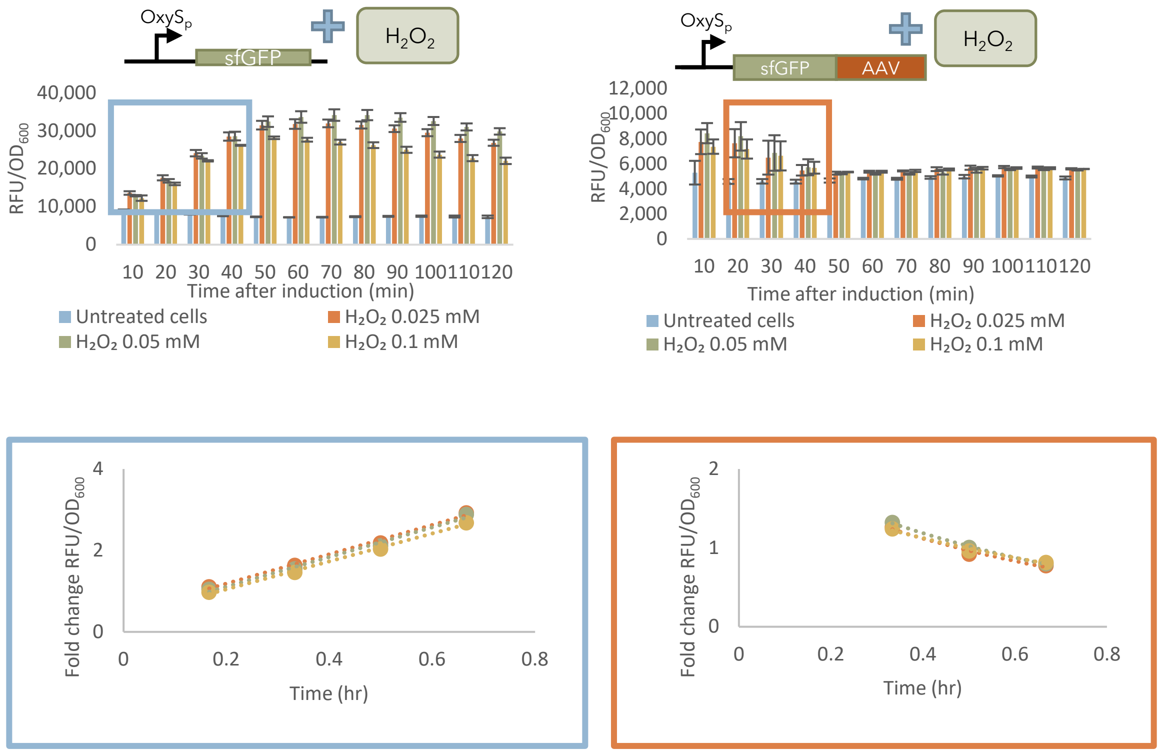


**Supplementary Figure S1** Figures **4a** and **4c** (top) are reproduced to highlight the regions used to calculate fluorophore expression and degradation rates (bottom).

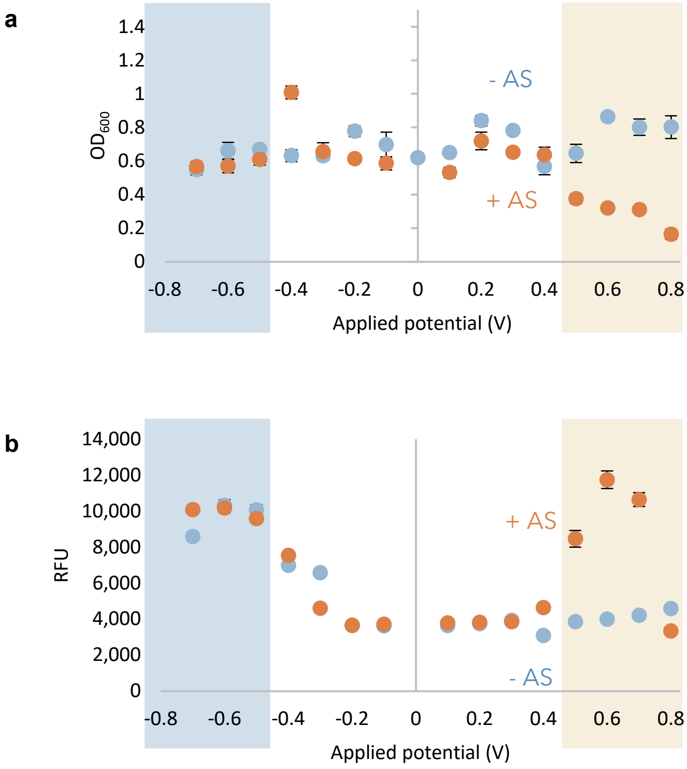


**<u>Supplementary Figure S2</u>** (**a**) $OD_{600}$ and (**b**) fold fluorescence of POxyRS-sfGFP reporter cells after varying potentials are applied for 7.5 minutes in the absence or presence of 500 μM AS. Values are taken after three hours of incubation.

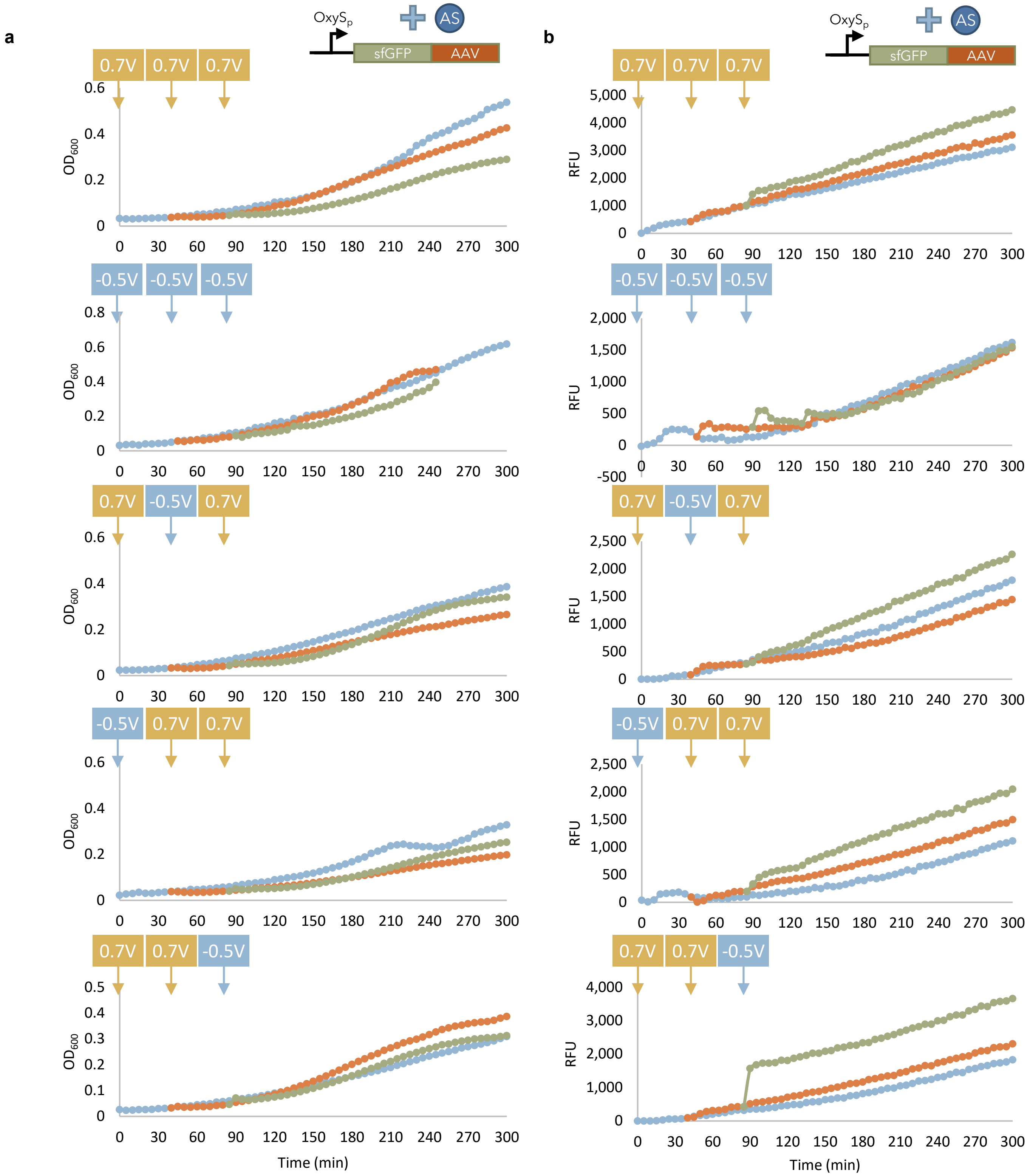


**Supplementary Figure S3** (**A**) $OD_{600}$ and (**B**) fluorescence of POxyRS-sfGFP-AAV reporter cells, in the presence of 500 μM AS, after application of "pulses" of potential at the indicated voltage and time for 7.5 minutes. If the value of fluorescence difference between the sample and the untreated cell negative control was less than 5 RFU, it was denoted as 0 RFU. Plots from top to bottom correspond to **Figures 7b-7f**.

**Supplementary Tables**

| Inducer | | sfGFP expression rate | sfGFP-AAV degradation rate |
|---|---|---|---|
| Inducer | Concentration (mM) | | |
| | | Zeroth-order | First-order |
| | | Units: ($RFU/OD_{600}$ fold change * $hr^{-1}$) ± standard error | Units: ($RFU/OD_{600}$ fold change * $hr^{-1}$) ± standard error |
| $H_2O_2$ | 0.025 | 3.598 ± 0.215 | -1.479 ± 0.254 |
| | 0.05 | 3.635 ± 0.261 | -1.515 ± 0.076 |
| | 0.1 | 3.415 ± 0.151 | -1.264 ± 0.152 |
| Oxidized AS | 0.25 | 1.007 ± 0.133 | 0.132 ± 0.059 |
| | 0.5 | 2.366 ± 0.407 | -0.021 ± 0.061 |
| | 0.75 | 3.864 ± 0.789 | -0.480 ± 0.048 |

**Supplementary Table S1** Expression (synthesis) and degradation rates of sfGFP expressed by OxyRS-sfGFP or OxyRS-sfGFP-AAV reporter cells treated with each inducer. Expression rates were calculated as a zeroth-order maximum rate from linear regression of sfGFP fluorescence (normalized to untreated as fold change $RFU/OD_{600}$). Based on the region of increasing fluorescence, sfGFP expression rates for $H_2O_2$-treated cells were calculated from 10 to 40 minutes after induction, while rates for oxidized AS-treated cells were calculated from 20 to 40 minutes after induction.. Degradation rates were calculated as a first-order decay rate from exponential regression ($b$ in $y = a * e^{bx}$) of sfGFP-AAV fluorescence (normalized to untreated as fold change $RFU/OD_{600}$). Based on the region of declining fluorescence, sfGFP-AAV degradation rates for $H_2O_2$-treated cells were calculated from 20 to 40 minutes after induction, while degradation rates for oxidized AS-treated cells were calculated from 70 to 100 minutes after induction. Standard errors of regression are reported.
